# Differential contribution of distinct neuronal populations to danger representations

**DOI:** 10.1101/2024.01.24.577067

**Authors:** Ana Paula Menegolla, Guillem Lopez-Fernandez, Cyril Herry, Mario Martin-Fernandez

**Affiliations:** Université de Bordeaux, Neurocentre Magendie, U1215, Bordeaux, France; INSERM, Neurocentre Magendie, U1215, Bordeaux, France

## Abstract

The recognition of specific stimuli and contexts in dangerous situations determines the expression of behaviors needed to appropriately cope with each threatening encounter. Moreover, the detection of common features shared by different dangerous situations allows eliciting general brain states and is necessary for both the expression of preparatory reactions and adaptive behavioral responses in a timely manner. However, it is unknown how general and specific danger representations emerge from the combined activity of different neuronal populations to elicit the expression of adaptive defensive responses. Using a behavioral paradigm that exposes mice to multiple threatening situations and calcium imaging recordings in freely moving mice, we investigated the role of different dmPFC neuronal populations in the generation of general and specific neuronal representations. Our results suggest that the population of somatostatin positive (SST^+^) interneurons generates specific representations while those arising from parvalbumin positive (PV^+^) interneurons are mainly unspecific. Together, this data suggests the presence of distinct information in different dmPFC neurons allowing a collective encoding of both general and specific danger representations.

## Introduction

The recognition of discrete features of stimuli and contexts informs about the specificities of animal surroundings. The activity of neuronal populations devoted to the formation of specific brain representations is essential to express optimal behavioral responses to properly cope with a particular threatening situation^1–4^. Moreover, the ability to detect and classify individual features into general categories is computationally advantageous and allows the generation of broader brain representations^5–7^ that are necessary for the expression of behaviors shared by distinct situations^3,4,8,9^. This is especially relevant when animals face threatening situations, as both preparatory reactions to danger and adaptive defensive responses to specific threats are essential to cope with the incoming danger. The survival of an organism hinges significantly on its capacity to recognize potential threats and to select suitable defensive responses to properly cope with danger. The expression of adapted defensive behaviors is key to help individuals avoid predation, escape from threats and protect themselves and their offspring. However, dysfunctions in the response to threats are implicated in various neurological and psychiatric conditions such as anxiety disorders, phobias, or post-traumatic stress disorder^10^. Thus, understanding the neural mechanisms underlying the expression of defensive behaviors in dangerous situations is essential to unravel the neuronal bases of these severe conditions^11–16^

In order to select the most advantageous defensive response, animals must be able to detect threatening situations and to discriminate the type of threat that is confronted. To achieve this, multiple interconnected brain regions shape survival neuronal circuits^17,18^ to identify the presence of a dangerous situation and accurately recognize the specificity of the threat encountered. Traditional fear-conditioning and active avoidance paradigms have been widely used to study the brain mechanisms underlying emotional processing of threat-related information^11,17,19–25^. However, these paradigms entail a reductionist design in which a single threat is presented and a single defensive response is examined. This has long been a limitation when trying to disentangle whether neuronal populations recorded in specific brain regions encode information specific to each threatening situation and/or more general information relative to the presence of a dangerous stimulus and the defensive state elicited. Nevertheless, a new generation of behavioral paradigms has been developed in which mice are not exposed to a single threat but also to multiple threats^1,26^ and/or stimuli of positive valence^1,27–33^. Similarly, we recently developed a behavioral paradigm, the Multi input-output (MIO) paradigm, in which mice are presented with multiple threats and elicit different defensive behaviors^3^. Using this task while performing electrophysiological recordings in the dorsal division of the medial prefrontal cortex (dmPFC), which has largely been evidenced as a key regulator of defensive behavior^13,14,23,27,34^, we showed that the dmPFC simultaneously encodes a general danger representation but also specific information about the identity of each threatening situation^3^.

Although the PFC contains distinct classes of neurons with different functional properties^35^, most of the literature on the study of defensive behaviors has either employed mixed populations or focused on the role of the most abundant neurons in the PFC, the excitatory pyramidal neurons^3,34,36,37^. Nonetheless, the activity of pyramidal neurons (Pyr) is extensively orchestrated by a network of GABAergic interneurons (INs) with diverse morphology, firing properties, postsynaptic targets and protein expression patterns^35,38^. Indeed, it is known that dmPFC processing of emotionally relevant information to control the expression of defensive behaviors does not solely depend on Pyr neuronal activity, but on the interaction between Pyr neurons and a heterogeneous set of interneuron populations^23,39–42^. Parvalbumin-expressing (PV^+^) interneurons, the most abundant of these interneuron types, mostly convey feedforward inhibition onto the soma and axon initial segment of Pyr neurons^35,38^. Somatostatin-expressing (SST^+^) interneurons are less abundant and mainly deliver feedback inhibition targeting the apical dendrites of excitatory Pyr neurons^35,38^. There is a growing body of evidence showing that, within the dmPFC, these two IN populations play a critical role in the expression of defensive behaviors^23,40–42^. However, it is still unclear how dmPFC Pyr neurons, SST^+^ and PV^+^ interneurons cooperate to generate general and specific representations to control the expression of defensive behaviors. To investigate the specific role of dmPFC neuronal populations in the encoding of general and specific representations of danger, we used calcium imaging to record the activity of dmPFC Pyr neurons, SST^+^ and PV^+^ interneurons populations in mice facing different threatening situations in the MIO paradigm. Our results suggest that although all the studied neuronal populations are modulated by the presence of threatening situations, each population of interneurons contributes to general and specific encoding of information differently. The activity of the population of SST^+^ interneurons allowed us to specifically discriminate between threatening or non-threatening trials as well as between the sensory inputs setting each threatening situation apart, while the population of PV^+^ interneurons encoded, in a more unspecific manner, the general presence of stimuli independently of their emotional value, the tone or the context.

## Results

To investigate the encoding of general and specific danger information within different dmPFC neuronal populations we used the multi input-output (MIO) paradigm^3^, a behavioral task that allowed us to study the neuronal activity of freely moving mice facing different threatening situations within the same behavioral session (**Fig. 1**). To monitor the activity of distinct neuronal populations within the dmPFC of mice performing the MIO paradigm we expressed the calcium indicator GCaMP6f in either CaMKII^+^, SST^+^ or PV^+^ neurons. We implanted gradient-index (GRIN) lenses in the dmPFC and used miniscopes (Inscopix) to image the calcium levels of the different neuronal types (**Fig. 1b**). To monitor the calcium levels of CaMKII-expressing pyramidal neurons we employed an AAV5-CaMKII-GCaMP6f viral construct in C57/BL6 mice. To image the calcium levels of SST-expressing and PV-expressing interneurons we used a Cre-dependent GCaMP6f viral construct (AAV5-Syn-Flex-GCaMP6f) in mice lines expressing Cre in SST^+^ or PV^+^ neurons respectively. These mice were then trained in the MIO paradigm. In this paradigm mice are freely exploring a rectangular behavioral box divided in 2 parts, the shelter and the arena (**Fig. 1a**). During active trials a conditioned stimulus (a constant tone of 7 kHz for a maximum of 7 seconds, the CS^+^1) is presented when mice are in the arena and it is associated with a mild foot shock (the unconditioned stimulus; US) that can be avoided by shuttling to the shelter. In contrast, during the passive trials the CS^+^1 is presented when mice are in the shelter and mice must stay in the shelter for the whole duration of the CS (7 seconds), otherwise they receive a foot shock if shuttling to the arena. During the inverse trials mice have to shuttle to the arena to avoid the US in response to a different tone (a constant tone of 2 kHz for a maximum of 7 seconds, the CS^+^2). Altogether, mice are presented with 3 different threatening trials requiring the integration of both tone and context to elicit the correct defensive responses in each threatening situation (**Fig. 1c**). Mice are also presented with control trials (a constant white-noise tone lasting 7 seconds, the CS^−^) that are never associated with a foot shock independently of mice location or behavior. Overall, mice learned to perform the MIO paradigm, eliciting the correct defensive responses in each threatening situation (**Fig. 1c**). The dmPFC neuronal activity of mice was imaged during the different training sessions and the day of best performance for each mouse was used to evaluate the activity of the different neuronal types during each trial type (**Fig. 1d**). As observed by the representative field of views (FOVs) of **Fig. 1d**, imaging pyramidal neurons gave a higher yield (∼140 neurons per FOV) than imaging SST^+^ and PV^+^ interneurons (∼35 and 25 neurons per FOV respectively). The higher number of pyramidal neurons within the imaged FOVs is consistent with the higher proportions of this neuronal type within the PFC^35^. Using this method we could then image the activity of pyramidal, SST^+^ and PV^+^ neurons (**Fig 1d**). We observed that: (1) All neuronal populations studied contained neurons responsive to CS^+^ and control trials, (2) single neuronal responses presented strong variability and that (3) response profiles varied greatly between neurons. These features are displayed in the 3 representative neurons shown in **Fig. 1d**. Together this data shows that the use of the recently developed MIO paradigm coupled with in vivo calcium imaging allows us to investigate the activity of different dmPFC neuronal types during multiple threatening situations and, therefore, the encoding of information related to either general or specific danger in pyramidal neurons, SST^+^ interneurons and PV^+^ interneurons.

**Figure 1.**
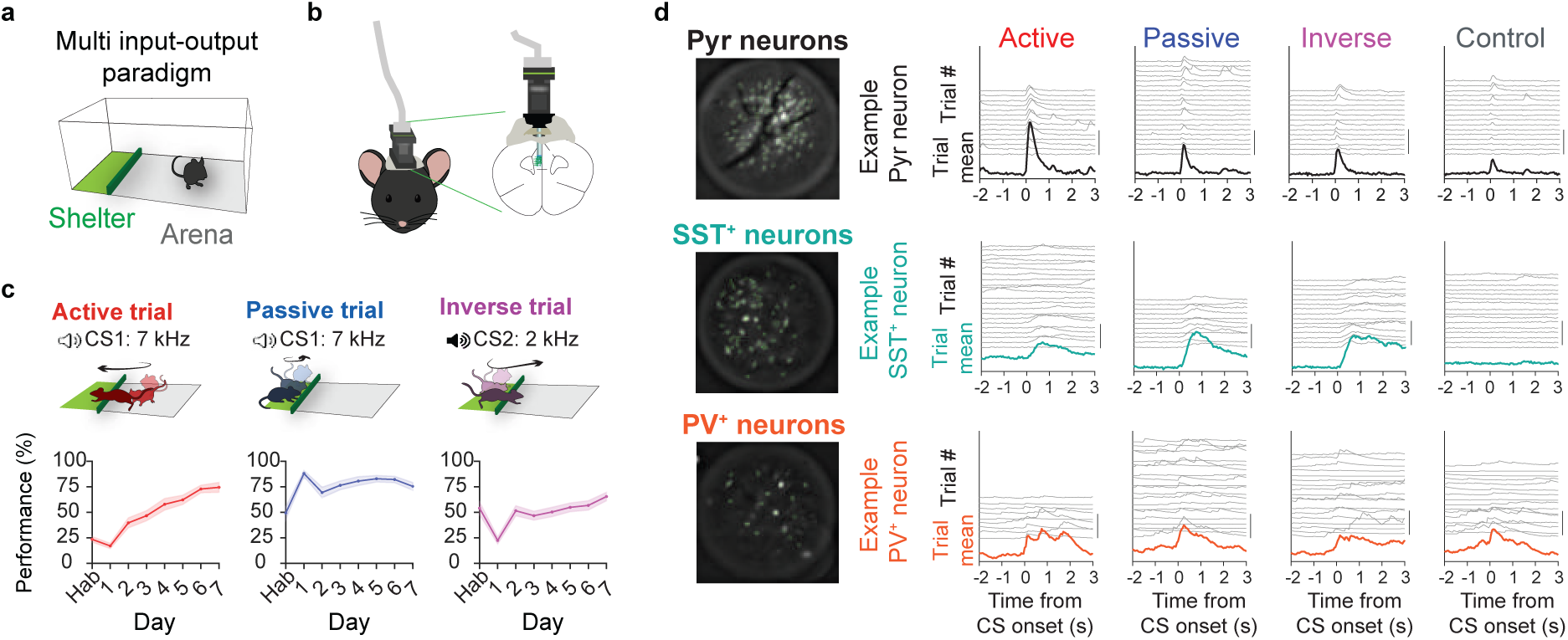
Calcium imaging recordings of Pyr, SST^+^ and PV^+^ neurons during the multi input-output paradigm. **a,** Representation of the behavioral box employed in the multi input-output (MIO) paradigm. The behavioral box is divided in shelter (green) and arena (grey). **b,** Schematic drawing showing the miniaturized endoscopes used to record the calcium levels of the different neuronal types in freely moving mice. **c, Top:** Schematic representations of the three threatening situations presented in the MIO paradigm (active, passive and inverse trials). **Bottom:** Average performance across mice in the MIO paradigm (n=23 mice) in active (red), passive (blue) and inverse (magenta) trials. Shaded areas represent S.E.M. **d, Left:** Representative fields of view of three mice showing the regions of interest detected with CNMFe containing pyramidal (black), SST^+^ (turquoise) and PV^+^ (orange) neurons. Fields of view obtained with lenses of 500 µm diameter. **Right:** Individual (grey) and mean (black, turquoise or orange) calcium traces of a pyramidal, SST^+^ and PV^+^ example neurons during active, passive, inverse and control trials. Note the variability and heterogeneity of neuronal responses. Scale bar 100% dF/F for single trial traces and 10% dF/F for mean trials.

To investigate responses of the different neuronal types during the different trials we looked at the activity of all the recorded neurons. We imaged 1253 pyramidal neurons from 9 mice, 211 PV^+^ neurons from 8 mice and 208 SST^+^ neurons from 6 mice (**Fig. 2**). We used this data to investigate the modulation of the different neuronal populations during the CS onset and the CS offset of active, passive, inverse and control trials (**Fig. 2a**). Aligning the neuronal activity to the CS onset allowed us to investigate the engagement of each neuronal populations at the onset of the threatening and control trials, while aligning their activity at the CS offset allowed us to investigate the neuronal modulation related to the expression of the different defensive behaviors (**Fig. 2**). Pyramidal neurons showed a clear increase in activated and inhibited neurons at the CS onset during threatening trials and also during control trials (although with briefer responses in control than in threatening trials; **Fig. 2b**). We also assessed the modulation of pyramidal neurons before and after the CS offset of the 3 threatening trials (active, passive and inverse) but not at the offset of the control trials (**Fig. 2b**), consistent with the lack of defensive response elicited during these trials. Then, we investigated the neuronal activity of PV^+^ interneurons during the different trial types. Similarly to pyramidal neurons, PV^+^ interneurons were activated and inhibited at the onset of the CS during all trial types. However, the responses of PV^+^ neurons seemed more sustained and appeared before and after the CS offset for all trial types (including control trials; **Fig. 2c**). Finally, SST^+^ interneurons showed response profiles strikingly different to those observed in pyramidal and PV^+^ interneurons (**Fig. 2d**). The population of SST^+^ interneurons contained a higher proportion of activated neurons when compared to the other investigated neuronal types. SST^+^ neurons showed stronger and more sustained modulations both at the CS onset and before and after the CS offset (**Fig. 2d and e**). Together, this data shows that all studied neuronal types are modulated (both through activation and inhibition) during threatening trials, suggesting that pyramidal neurons, SST^+^ and PV^+^ interneurons participate in the processing of threatening information in the dmPFC. Interestingly, CS^+^ trials strongly affected the activity of SST^+^ interneurons, suggesting a greater engagement of this neuronal population in the processing of information related to threatening trials than the other investigated neuronal types. Whereas this data shows that all the studied neuronal populations are modulated during threatening trials, it does not address the question of whether their activity is devoted to the encoding of information related to either general or specific danger.

**Figure 2.**
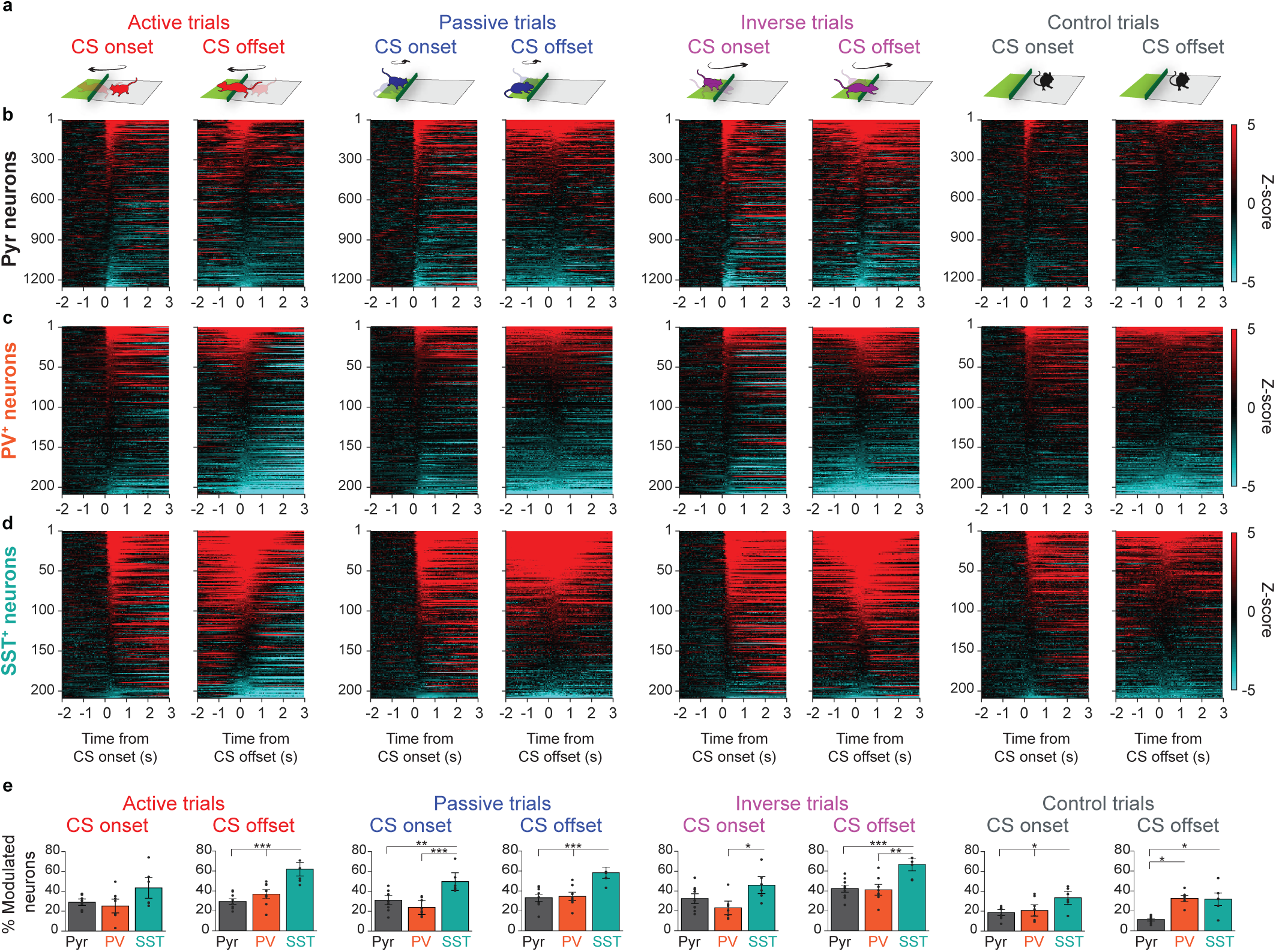
Activity of Pyr, SST^+^ and PV^+^ neurons during CS onset and offset. **a,** Schematic representations of mice behavior during CS onset and CS offset of active, passive, inverse and control trials. **b,** Z-scored mean activity of pyramidal neurons (black, n=1253 neurons, from n=9 mice) during CS onset (left) and CS offset (right) of active, passive, inverse and control trials. **c,** As **b,** but for PV^+^ neurons (orange, n=211 neurons, from n=8 mice). **d,** As **b,** but for SST^+^ neurons (turquoise, n=208 neurons, from n=6 mice). **e,** Mean percentage of modulated (activated + inhibited) neurons at CS onset (from 0 to 1 seconds after CS onset) and CS offset (from 0 to 1 seconds from CS offset) across mice in active, passive, inverse and control trials (n=9, 8 and 6 mice for Pyr, PV^+^ and SST^+^ neurons respectively). Small points represent percentages of modulated neurons in individual mice. Error bars represent S.E.M.

In order to address this, we used a population decoding approach based on linear kernel SVM classifiers applied to the neuronal activity of the different populations of neurons (See Methods). This strategy is illustrated in **Fig. 3a**, in which the calcium levels from the ensemble of neurons of each type of the recorded populations at a given time point t (the so-called population vector) is compared between conditions. In the scheme, active and control trials are represented; we repeated the same strategy between the following conditions: active, passive, inverse, control and no-CS trials (trials lacking any tone presentation). To obtain results comparable between the different neuronal populations we made classifiers using 100 randomly selected neurons of each type (**Fig. 3a**). Using this approach we were able to significantly decode with high decoding accuracies all threatening trials (active, passive and inverse trials) from the no-CS trials using either the population of pyramidal neurons or the populations of SST^+^ or PV^+^ interneurons (**Fig. 3b**). Moreover, we addressed whether using this approach we could decode the threatening trials from the control trials. Interestingly, we observed high and significant decoding accuracies at CS onset when the populations of pyramidal neurons and SST^+^ interneurons were challenged to classify threatening from control trials. However, the population of PV^+^ interneurons presented only low decoding accuracies when decoding control from threatening trials (**Fig. 3b**). To investigate the number of neurons needed by the different neuronal populations to decode the threatening trials from either basal conditions (no-CS) or control trials, we constructed classifiers with a variable number of randomly selected neurons of each type (from 1 to 200 neurons of each type) and calculated the mean decoding accuracies obtained at CS onset (**Fig. 3c**). Using this technique we observed that SST^+^ neurons provided higher decoding accuracies even when the classifiers used lower number of neurons. Moreover, while the population of pyramidal neurons and SST^+^ interneurons obtained similar decoding accuracies when decoding threatening trials from either control trials or trials with no tone presentation (no-CS trials; **Fig. 3c**), the population of PV^+^ interneurons showed a clear decrease in the decoding accuracies when decoding threatening from control trials as compared to threatening vs no-CS trials (**Fig 3c**). This indicated that, although PV^+^ interneurons could decode the presence of a tone from baseline, their population activity did not allow us to decode whether the tone presented was threatening or not (CS^+^ or control tone; unlike SST^+^ and pyramidal neurons. **Fig. 3b and c**). This suggests that the population of PV^+^ interneurons carried more unspecific information than SST^+^ and pyramidal neurons. To further confirm this observation, we tested the generalization of the classifiers among threatening and control trials (see Methods). To investigate the threat generalization, we trained a classifier to decode a random threat type (active, passive or inverse) from basal activity (no-CS trials) and challenged that classifier to classify a different threat type (not used to train the classifier) from basal activity. Using this approach, we observed that the classifiers made with all the studied neuronal populations showed high levels of generalization across the threatening trials (Threat-generalization; **Fig. 3d**). Then we investigated whether these neuronal populations could decode control trials from no-CS trials and observed a transient decoding accuracy (around 500 ms) in the pyramidal and PV^+^ neuronal populations and a sustained encoding in the SST^+^ interneuronal population (**Fig. 3d**). Next, we evaluated the generalization of the patterns of activity presented in the threatening trials to the control trials. For this, we used the same classifier that we trained for the threat generalization to classify control trials from basal activity. We observed that, although SST^+^ interneurons showed the strongest encoding of control trials, they lacked strong levels of generalization from threatening to control trials (**Fig. 3d**). The SST^+^ and the pyramidal neuronal populations only presented a transient control-generalization while PV^+^ neuronal population showed levels of control-generalization similar to those obtained when decoding control trials (**Fig. 3d**). We then measured the accuracy loss as the difference in accuracy obtained in each iteration between the ‘threat’ classification accuracy and the ‘threat-general’ (CS^+^-CS^+^) and ‘control-general’ (CS^+^-control) classification accuracies. Indeed, there was a stronger accuracy loss in control-generalization than in threat-generalization for pyramidal and SST^+^ neuronal populations, while this difference was not present in the PV^+^ neuronal population (**Fig. 3e**). These results show that the patterns of activity presented in the population of PV^+^ interneurons at the early CS onset (500 ms) are shared among the different trials, independently of whether those are threatening or not. This data, together with the low decoding accuracies obtained between threatening and control trials (**Fig. 3b and c**) suggests that the population of PV^+^ interneurons encodes the presence of trials (or auditory stimuli) in an unspecific manner. However, the population of SST^+^ interneurons showed stronger decoding accuracies than the other studied neuronal populations when decoding threatening from no-CS trials (**Fig. 3b and c**), threatening from control trials (**Fig. 3 b and c**) and control from no-CS trials (**Fig. 3d**), with low generalization between threatening and control trials (**Fig. 3d and e**). This points to the encoding of information by SST^+^ populations being more robust than the encoding present in PV^+^ and pyramidal neurons in both threatening and non-threatening trials and that this is achieved with different patterns of activity dependent on the emotional value of the stimuli. We then investigated whether the activity of the different neuronal populations encoded information necessary for the correct identification of the threatening trials and the selection of defensive responses. To do so, we took advantage of the design of the behavioral paradigm, in which mice need to discriminate both tone and context to select the correct defensive response in each threatening situation (**Fig. 1c**). Then we evaluated whether the dmPFC population activity could decode threatening trials that presented different tones (CS^+^1: active and passive trials; CS^+^2: inverse trials) and threatening trials in different contexts (Shelter: inverse and passive trials; Arena: active trials) and observed that in both conditions the population activity of SST^+^ interneurons provided stronger decoding accuracies than the populations of pyramidal and PV^+^ neurons (**Fig 3f and g**). This suggests that the population of SST^+^ interneurons provides not only a stronger encoding of threatening and non-threatening trials but also more specific information about the different threatening trials presented. Together, this data suggests that, although all the studied neuronal populations are modulated during the threatening trials, their activity encodes different information. While the pyramidal population activity encodes the presence of threatening trials specifically, the population of PV^+^ interneurons encodes the presence of stimuli independently of their emotional value, and the population of SST^+^ interneurons not only informs about the threatening or non-threatening nature of the trials, but also provides specific information about the different factors differentiating the threatening trials (tone and context).

**Figure 3.**
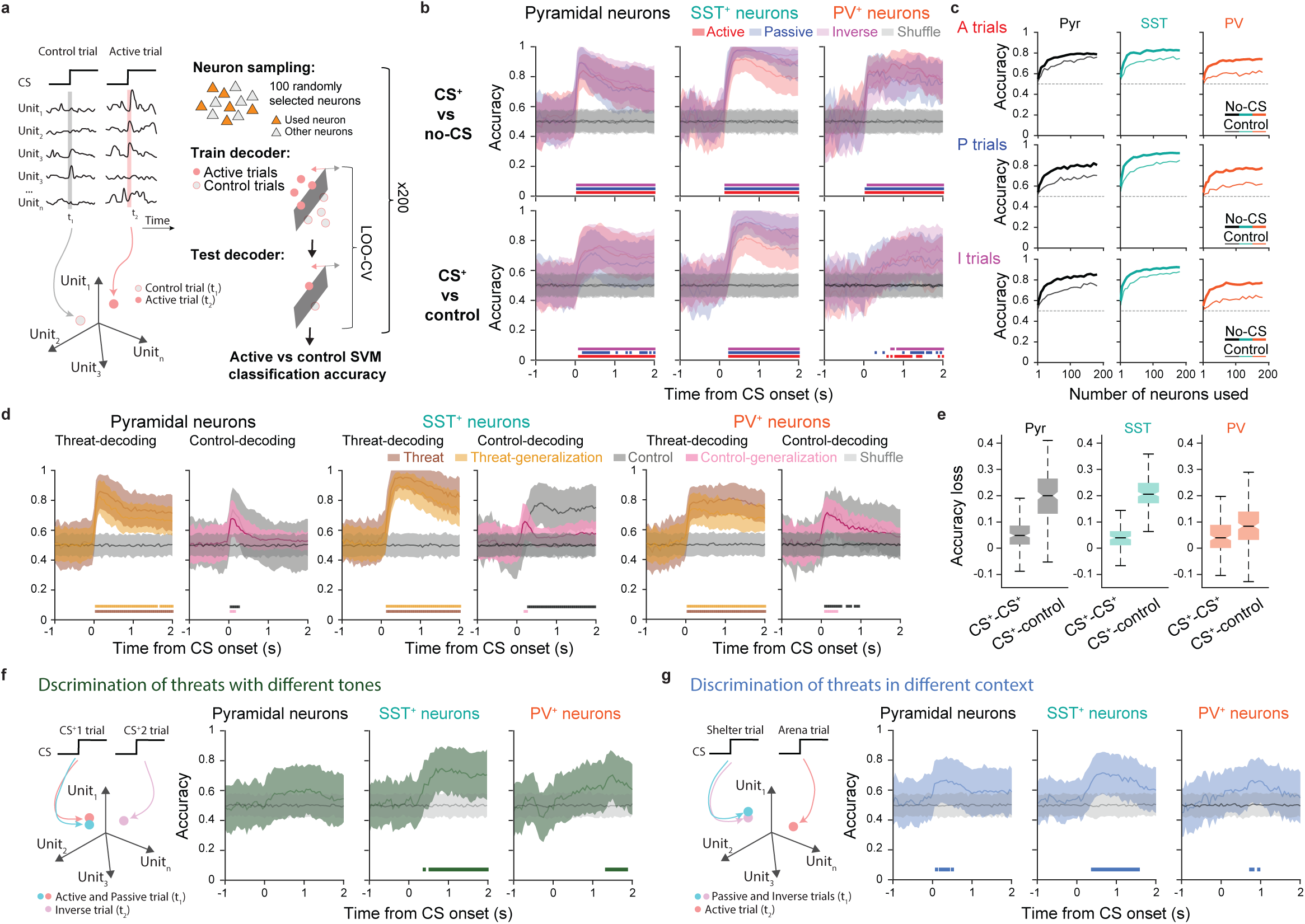
Encoding of threat-related information in different dmPFC neuronal populations. **a,** Schematic representation of the strategy employed to construct the pseudo-population vectors and the support vector machine (SVM) classifiers used to decode the patterns of population activity presented in the different trials. **b, Top:** Classification accuracies obtained around CS onset for classifiers challenged to decode between CS^+^ trials and no-CS conditions during active (red), passive (blue) and inverse (magenta) trials. Shuffle decoding accuracies in grey. Significant classification accuracies represented as horizontal bars in red (active trials), blue (passive trials) and magenta (inverse trials). **Bottom:** As top but showing classification accuracies obtained for classifiers challenged to decode between CS^+^ and control (CS^−^) trials. **c,** Mean classification accuracy between CS^+^ and no-CS trials (thick lines) and CS^+^ and control trials (thin lines) obtained with classifiers that used different numbers of neurons. Classifiers used active (top), passive (middle) and inverse (bottom) trials and Pyr (black), SST^+^ (turquoise) and PV^+^ (orange) neurons. **d,** Average threat (brown), threat-general (orange), control-general (pink) and control classification accuracies for classifiers built with Pyr, SST^+^ and PV^+^ neurons. Shuffle decoding accuracies in grey. Significant classification accuracies represented as horizontal bars in brown (threat-decoding), orange (threat-general), pink (control-general) and black (control). **e,** CS^+^-CS^+^ and CS^+^-control accuracy loss (see Methods) in Pyr, SST^+^ and PV^+^ neurons. Box plot center line represents median; box limits are 25th and 75th percentiles; whiskers are more extreme non-outliers data points. **f, Left:** Schematic representation of the strategy employed to decode among threats with different tones. **Right:** Tone-decoding accuracies relative to the CS^+^ onset in Pyr, SST^+^ and PV^+^ neurons. **g,** As **f,** but for the decoding of threats presented in different contexts.

## Discussion

Coupling the recently developed MIO behavioral paradigm with calcium imaging in freely moving mice we were able to investigate the generalization and specificity of neuronal representations in different neuronal populations. Our results suggest that different neuronal types represent different information allowing the dmPFC to encode both general and specific danger.

The MIO paradigm was previously developed to investigate the neuronal activity of mice facing different threatening situations during the same behavioral session^3^. This task offers an advantage from classical behavioral paradigms, in which mice are presented with a single threatening event, since it permits us to evaluate whether the neuronal responses evoked by a threat are either specific or general for all dangerous situations. We coupled this paradigm with the imaging of calcium activity of different dmPFC neuronal types in freely-moving mice using miniscopes (**Fig. 1**). We imaged the activity of Pyr neurons, SST^+^ and PV^+^ interneurons, generating the first dataset in which large populations of dmPFC SST^+^ and PV^+^ interneurons are imaged in mice performing defensive behaviors. Moreover, coupling this paradigm with calcium imaging, this dataset allowed us to further investigate the specificity/generality of the CS^+^-evoked neuronal responses in distinct neuronal populations.

To investigate this, we first evaluated the responsiveness of each neuronal type during threat presentation and observed that all of them were modulated by the presence of threatening trials (**Fig. 2**). This suggests that all the studied neuronal populations are involved in the processing of danger-related information in the dmPFC. However, different neuronal populations showed different profiles of responsiveness. The most striking difference attains the SST^+^ interneurons: a higher percentage of this neuronal population was modulated from the onset to the offset of both threatening and control trials (**Fig. 2**). This suggests that SST^+^ interneurons display a higher degree of engagement in the processing of danger than either PV^+^ interneurons or pyramidal neurons. This could be due to their role in the dmPFC micro-circuitry^35^ or to their higher sparsity in the dmPFC. To further clarify the nature of this higher recruitment of SST^+^ interneurons further studies investigating the functional interactions of the different neuronal types in the dmPFC micro-circuitry during threat presentations are needed. However, despite the higher modulation of SST^+^ interneurons during threatening trials, all neuronal types showed evoked responses to threat during all trial types.

To investigate whether the activation of the different neuronal types was either specific for each threatening situation or more general, we looked at the population patterns of activity elicited by the different trials in each neuronal population. Although the presence of threatening trials could be decoded by the population activity of all neuronal types, the activity of PV^+^ interneurons provided unspecific information that did not allow us to differentiate between threatening and non-threatening (control) trials (**Fig. 3**). In strike contrast, the population of SST^+^ interneurons displayed patterns of activity that, not only allowed us to discriminate between threatening and non-threatening trials, but also between threatening trials presenting different tones (CS^+^1 or CS^+^2) or contexts (shelter or arena; **Fig. 3**). Pyr neurons displayed patterns of activity that allowed us to discriminate between threatening and non-threatening situations but that did not allow us to classify threatening trials presenting different features. Although this result is consistent with the primacy of general representations of danger observed in the mixed population of dmPFC^3^, it is important to note that the presented results are obtained sampling an equal number of neurons for each population (100 neurons). An alternative approach would be to evaluate the classification accuracies adjusting the numbers of neurons used for each population in a manner that resembled the proportions observed in the dmPFC (approximately 80% Pyr neurons, and 20% interneurons). Therefore, the classification accuracies obtained with a larger population of Pyr neurons may serve as a better comparison for those obtained with 100 SST^+^ and PV^+^ interneurons. Nonetheless, our results show a differential encoding of general and specific representations of danger in the dmPFC interneuron populations. SST^+^ interneurons show more specific representations and their population activity allowed us to classify not only the threatening nature of the stimuli, but also the sensory features distinguishing the threatening trials (tone and context). These results suggest that these neurons may provide specific inhibition necessary for the appearance of specific threat-representations in the dmPFC. Moreover, this data further suggests that SST^+^ activity is necessary for the discrimination of threatening tones and contexts, however, further studies demonstrating the causality of the involvement of SST^+^ interneurons in the selection of defensive responses are needed. The PV^+^ population activity was largely unspecific and did not allow us to discriminate between control and threatening trials. Further studies are needed to disambiguate whether their activity in threatening situations is passively informing about the presence of an auditory stimuli or whether it promotes the engagement of mice in the behavioral task by favoring processes such as arousal or attention.

Overall, our study shows a distinct role of different interneuron populations in the processing of threat-related information within the prefrontal cortex. Maladaptive fear processing may occur both through the generalization of danger-representations to non-threatening situations or through the impairment of the threat-specific representation, which could affect the ability to adaptively cope with threats. A better understanding of the processes that allow the dmPFC to generate both general-danger and specific-threat representations might shed light in the mechanisms responsible of the development of those pathological conditions.

## Materials and Methods

### Subject detail

Male C57BL6/J (8-24 weeks old, Janvier), SST-IRES-Cre (8-24 weeks old, Jackson Laboratory, Ssttm2.1(cre)Zjh/J) and PV-IRES-Cre (8-24 weeks old, Jackson Laboratory, B6;129P2-Pvalbtm1(cre)Arbr/J) mice were single-housed for at least three weeks before experimentation, under a 12-h light-dark cycle, and provided with food and water *ad libitum*. The housing temperature was 22 ± 1°C, and housing humidity was 60% ± 5%. All procedures were performed per standard ethical guidelines (European Communities Directive 86/60-EEC) and were approved by the Animal Health and Care committee of Institut National de la Santé et de la Recherche Médicale and French Ministry of Agriculture and Forestry (authorization #A3312001). Mice were handled and habituated to be connected for three days before the experiment started. All data correspond to implanted and connected animals.

### Behavior

Experiments were run in a behavioral box (40 x 10 x 30 cm) composed of Plexiglas walls and a grid floor used to deliver footshocks (0.6 mA) located in a sound-attenuating chamber. This box was divided in the shelter zone (5 x 10 cm) and the arena zone (35 x 10 cm) by a small plastic hurdle (0.5 cm high). The shelter zone was marked with visual cues to facilitate its recognition. The box was equipped with infrared beams to detect mice shuttling between the shelter and arena zones. The behavioral box was enclosed in an acoustic isolated box containing speakers, a light source and a video camera (30 frames per second; Cineplex, Plexon) above the behavioral box. Three different auditory cues were delivered at a volume of 75 dB: CS^+^1 (7 kHz), CS^+^2 (2 kHz) and CS^−^ (white noise). The auditory cues were presented for 7 s or until mice shuttled from one zone to the other (shelter and arena) in the case of the CS^+^. The CS^−^ was presented for 7 s independently of mouse behavior.

The presence of the auditory cue and the location of the mice in either the shelter or the arena established the different trial types. In active trials, a CS^+^1 was presented when mice were in the arena. Mice had to shuttle to the shelter before the end of the CS (7 s) to avoid a foot shock (1 s). In passive trials, the CS^+^1 was presented when mice were in the shelter. Mice had to remain in the shelter for the duration of the CS (7 s) to avoid a foot shock (1 s). In inverse trials, a CS^+^2 tone was presented when mice were in the shelter. Mice had to shuttle to the arena before the end of the CS (7 s) to avoid a foot shock (1 s). In control trials, the CS^−^ was presented either in the shelter or in the arena without any reinforcement and independently of mouse behavior. The foot shocks lasted for a maximum of 1 s (shuttling to the correct area of the box would end the foot shock) and co-terminated with the CS when mice did not perform the correct defensive response for each CS^+^ trial type (error trials). Therefore, at the end of error trials, there was an extra second of CS^+^–US co-presented. The different trial types were intermingled and dependent on mouse location and tone presentations. In a behavioral training session, mice were exposed to 64 intermingled and pseudorandomized ‘tone-presentation events’ of three different types: 24 type A (CS^+^1 presentations: active or passive trials according to mouse location), 24 type B (CS^+^1 presentation if mice were in the arena and CS^+^2 if mice were in the shelter: active or inverse trials, respectively) and 16 control (CS^−^ presentation independent of mouse location). Additionally, mice were exposed to 8 intermingled control trials with a duration of 7 seconds in which no tone was presented (no-CS trials). The ITI was pseudorandomized, ranging from 32 to 75 s. A behavioral session therefore lasted approximately 75 min on average. The training sessions consisted of a habituation session (identical to a normal training session but lacking shock presentations) and a maximum of nine training sessions until mice reached high behavioral performance levels for the three threatening trial types (active, passive and inverse trials). We calculated the performance of active, passive and inverse trials as the number of correct trials over the total number of trials for each trial type. The experimental box was cleaned with 70% ethanol before and after each session.

### Surgical procedure

Mice (8-10 weeks old) were anesthetized with isoflurane (induction 3%, maintenance 1.75%) in O2. Body temperature was maintained at 37 °C with a Temperature Controller System (FHC), and eyes were hydrated with Ocry-gel (TVM). For analgesia, a subcutaneous injection of 0.05 mL of Metacam (5 mg/kg body weight) was administered 30 min before anesthesia. Additionally, 0.1 mL of local lidocaine anesthesia (Lidor, 20 mg/mL diluted with sodium chloride at 0.5%) was applied under the scalp before incision. Mice were placed in a stereotaxic frame (Kopf instruments), and three stainless steel screws were attached to the skull before craniotomy to secure all implants.

### Virus injection and GRIN lens implantation

For GCaMP6f expression we used 280 nL of the following GCamP6f viral constructs: For CamKII^+^ neurons we used the AAV5.CamKII.GCaMP6f.WPRE.SV40 virus, titer 2.3×10^13^, Adgene 100834, injected in C57BL6/J mice; For SST^+^ or PV^+^ neurons we used the AAV5.Syn.Flex.GCaMP6f.WPRE.SV40 virus, Adgene 100833, titer 3.81×10^13^, in SST-IRES-Cre mice and PV-IRES-Cre mice, respectively). Following craniotomy, the corresponding viral solution was unilaterally injected into the dmPFC (+2.15 mm AP, ±0.35 mm ML relative to the bregma, and −1.4 mm DV from the dura) using a micromanipulator (Scientifica) and pulled glass pipettes (tip diameter ∼25 mm). After injection, a track above the imaging site was opened with a sterile needle (26G, 0.45 mm outer diameter, Terumo) to assist in inserting the GRIN lens (Gradient-index optics lens, 4 mm long x 0.5 mm wide). The needle was inserted at the exact coordinates of the injection, except for the DV, at 1.0 mm. The GRIN lens was then implanted at the following coordinates: +2.15 mm AP, ±0.35 mm ML relative to the bregma and −1.25 mm DV from the dura. Super-Bond cement was used to secure the GRIN lens. The upper surface of the skull was protected with Kwik-Sil silicone adhesive (World Precision Instruments).

### Calcium imaging recordings

Two weeks after surgery, to allow for optimal GCaMP6f expression, a miniaturized fluorescence endoscope (miniscope) baseplate was attached to the implant in anesthetized mice. The miniscope was used for GCaMP6f fluorescence detection so that the baseplate was fixed to the implant based on an optimal field of view within the dmPFC. The miniscope was detached, and the baseplate was covered with a cover. Experiments did not start earlier than 2 weeks after baseplating to allow further viral expression and recovery. One week before starting the behavioral training, mice were habituated to the mounting procedure and the weight of the miniscope for four consecutive days. For the first three days, animals were restrained, the baseplate cover removed, a miniscope dummy was attached to the baseplate, and mice were left with it for at least 30 minutes in their home cage. On the fourth day, mice were mounted with the miniaturized endoscope to set the miniature endoscope light-emitting diode (LED) power and electronic focus. Imaging data were acquired using the nVoke software at a frame rate of 20 Hz (exposure time, 50 ms) with a LED power from 1 to 1.5 mW/mm2, a gain from 1 to 3 and a spatial downsampling by a factor of 2. Recordings of calcium imaging activity during behavioral sessions were recorded in segments around the trial presentations. Imaging started 7 seconds prior to the onset of every trial (CS^+^, CS^−^ and no-CS trials), continued for the whole duration of the trial and ended 3 seconds after the end of the trial (7 seconds pre-trial + 1-8 seconds CS/US or no-CS + 3 seconds post-trial). Therefore, each recording segment lasts from 10 to 18 sec.

### Processing of calcium imaging data

The recordings were processed using the Inscopix Data Processing Software. The time series were joined, pre-processed, spatially filtered and motion corrected before identifying the cells in the field of view. Cell identification was performed through a CNMFe (*Constrained Nonnegative Matrix Factorization for microEndoscopic data; Zhou et al., 2018*) algorithm and each trace was validated or rejected by the experimenter based on the identified morphology, activity trace and calcium transients. We obtained the single-cell denoised calcium dF/F traces from each accepted neuron. Neural activity of all recorded mice was aligned to trial onset and the activity of each individual neuron was z-scored using a reference period of -2 to 0 seconds before CS onset. To quantify the percentage of modulated cells at CS onset and offset, we calculated the mean z-score activity aligned at CS onset (0 to 1 s from CS onset) or CS offset (0 to 1 s from CS offset) and considered individual neurons to be active or inhibited if their mean activity was above 1.96 or below -1.96, respectively. We then calculated the percentage of modulated cells as the total number of active and inhibited cells over the total number of recorded cells for each recorded mouse.

### Anatomical and histological analyses

Mice were euthanized with a solution containing pentobarbital (Exagon, 0.27 mg/g) and lidocaine (Lidor, 0.03 mg/g) and perfused via the left ventricle with 4% w/v paraformaldehyde (PFA) in 0.1 M PBS. Following dissection, brains were postfixed in PFA for 24h at 4 °C. Brain sections of 60-mm thickness were serially cut using a vibratome, mounted on gelatin-coated microscope slides and protected with vectashield antifade mounting medium (Vector Laboratories) and a coverslip. Slices were imaged using an epifluorescence system (Leica DM 5000) fitted with a 10x dry objective. The location of GRIN lens implantation and extent of the viral injections were visually controlled. Only infections accurately targeting the dmPFC and GRIN lenses terminating within this region were considered for the analyses.

### Population analyses and decoding

To investigate the activity of the population of recorded neurons, the calcium activity from the ensemble of recorded cells at a given time point *t* was pooled in a vector containing as many dimensions as neurons recorded. This instantaneous population vector contained the calcium activity of all the recorded neurons and mice at each time point. For population activity we used the neuronal activity analyzed in 50-ms time bins and relative to the 2 seconds prior to CS presentation by subtracting the mean dF/F in that time period from the trial. To assess the decoding accuracies of the dmPFC population, we used linear kernel SVM classifiers in single-trial instantaneous pseudo-population vectors (including units and trials from different mice) composed of the calcium activity of each neuron (50-ms time bins) after the onset of the five different trial types (active, passive, inverse, CS^−^ and no-CS trials). For the decoding of threat versus threat, threat versus control, threat versus no-CS or control versus no-CS, the decoding was performed among population vectors at time *t* in relation to the onset of two different trial types. To obtain comparable accuracies across the classifiers used in the different conditions, we used for all classifiers the same number of trials to train and test decoders. We used a leave-one-out cross-validation (LOO-CV) method from a total number of 5 trials for each class. To account for the different number of neurons in each dataset we subsampled 100 random neurons for each iteration of the classifier. Using this method, we first selected 100 random neurons and trained a classifier cross-validated by a LOO-CV method. To obtain the average decoding accuracies, this technique was repeated 200 times, randomly selecting in each iteration 100 different neurons and different trials to construct the single-trial pseudo-population vectors. We used 5 trials of each class for all classifiers to obtain comparable decoding accuracies among the different classifiers. To evaluate the difference between the obtained decoding accuracies with the accuracies that could be obtained by chance, we compared the decoding accuracies of our classifiers with that obtained using a ‘shuffled’ condition. To do this, we employed the same method described above but in which the class identity of the test dataset was randomly assigned. Next, we used a two-sided permutation test by computing P values as the proportion of the shuffle repetitions that exceeded the decoding accuracies obtained with the method above using the real identity of classes. Significant decoding accuracies were defined as P < 0.05. To investigate the generalization of the population activity patterns in the different neuronal types we used the previous method to train a classifier to decode a random threat type (active, passive or inverse) from basal activity (no-CS trials) and tested it with the same classes to obtain the ‘threat’ decoding accuracy. We then challenged that classifier to classify a different threat type (not used to train the classifier) from a no-CS trial, obtaining the ‘threat-general’ decoding accuracy. The same classifier was used to classify the control trials from the no-CS to obtain the ‘control-general’ classification accuracies. We measured the CS^+^-CS^+^ and CS^+^-control accuracy loss as the difference in accuracy obtained in each iteration between the ‘threat’ classification accuracy and the ‘threat-general’ and ‘control-general’ classification accuracies. For tone and context discrimination decoding, the same decoding strategy and statistical methods described above were used. For Tone Discrimination decoding, 5 randomly selected trials among active and passive trials were used as a single class decoded from 5 randomly selected inverse trial. For Context discrimination 5 randomly selected trials among inverse and passive trials were used as a single class decoded from 5 randomly selected active trial.

### Statistics and reproducibility

No statistical methods were used to predetermine sample size. Experiments were not randomized, and investigators were not blinded to allocation during experiments and outcome assessment. We conducted all analyses using either custom routines written in MATLAB (Mathworks) or SigmaPlot (Systat Software). Figures were assembled using Adobe Illustrator. For all datasets, normality was tested using Kolmogorov-Smirnov test (*α* < 0.05) and homogeneity of variance with Levene’s test (*α* < 0.05) to determine whether parametric or non-parametric analyses were required. We used parametric analyses including one-way ANOVA followed by post hoc Tukey’s test if a significant main effect or interaction was observed. For all tests, \**P* < 0.05, \*\**P* < 0.01, \*\*\**P* < 0.001 and not significant >0.5; significance was calculated after correction for multiple comparisons. If either homogeneity of variance or normality assumptions were not met, nonparametric analyses were used. When required, we used nonparametric Kruskal-Wallis one-way ANOVA on ranks and Dunn’s multiple-comparison post hoc test to protect from false positive errors. All behavioral and imaging data were collected in a computer-controlled, automated and unbiased manner. No statistical methods were used to predetermine sample sizes, but sample sizes were similar to those reported in previous publications^7,33^

## Author contribution

A.P.M., G.L.-F and M.M-F. performed calcium imaging recordings. A.P.M., and G.L.-F. performed histology. A.P.M., G.L.-F and M.M-F performed surgeries. M.M.-F. and C.H. designed experiments. M.M.-F. designed population analyses and wrote software codes. A.P.M., G.L.-F, C.H., and M.M-F. analyzed data and wrote the paper. All the authors read and edited the manuscript.

## Acknowledgments

We thank S. Laumond, J. Tessaire and the technical staff of the housing and experimental animal facility of the Neurocentre Magendie. This work was supported by EMBO (ALTF 200-2018) and HFSP postdoctoral fellowships (LT0000658/2019) to M.M.-F., ECO-Contrat doctoral de la Fondation pour la Recherche Médicale (ECO202206015556) to G.L.-F., grants from the French National Research Agency (ANR-10-EQPX-08 OPTOPATH), the Conseil Regional d’Aquitaine and the Fondation pour la Recherche Médicale (FRM-Equipes FRM 2017). The funders had no role in study design, data collection and analysis, decision to publish or preparation of the manuscript.

## Competing interests

The authors declare no competing interests

## Notes

### Competing Interest Statement

The authors have declared no competing interest.

